# *Aspergillus fumigatus* acetate utilisation impacts virulence traits and pathogenicity

**DOI:** 10.1101/2021.06.09.447827

**Authors:** Laure Nicolas Annick Ries, Patricia Alves de Castro, Lilian Pereira Silva, Clara Valero, Thaila Fernanda dos Reis, Raquel Saborano, Iola F. Duarte, Gabriela Felix Persinoti, Jacob L. Steenwyk, Antonis Rokas, Fausto Almeida, Jonas Henrique Costa, Taicia Fill, Sarah Sze Wah Wong, Vishukumar Aimanianda, Fernando José Santos Rodrigues, Relber A. Gonçales, Cláudio Duarte-Oliveira, Agostinho Carvalho, Gustavo H. Goldman

## Abstract

*Aspergillus fumigatus* is a major opportunistic fungal pathogen of immunocompromised and immunocompetent hosts. To successfully establish an infection, *A. fumigatus* needs to use host carbon sources, such as acetate, present in the body fluids and peripheral tissues. However, utilisation of acetate as a carbon source by fungi in the context of infection has not been investigated. This work shows that acetate is metabolised via different pathways in *A. fumigatus* and that acetate utilisation is under the regulatory control of a transcription factor (TF), FacB. *A. fumigatus* acetate utilisation is subject to carbon catabolite repression (CCR), although this is only partially dependent on the TF and main regulator of CCR CreA. The available extracellular carbon source, in this case glucose and acetate, significantly affected *A. fumigatus* virulence traits such as secondary metabolite secretion and cell wall composition, with the latter having consequences for resistance to oxidative stress, to anti-fungal drugs and to human neutrophil-mediated killing. Furthermore, deletion of *facB* significantly impaired the *in vivo* virulence of *A. fumigatus* in both insect and mammalian models of invasive aspergillosis. This is the first report on acetate utilisation in *A. fumigatus* and this work further highlights the importance of available host-specific carbon sources in shaping fungal virulence traits and subsequent disease outcome, and a potential target for the development of anti-fungal strategies.

**Importance:** *Aspergillus fumigatus* is an opportunistic fungal pathogen in humans. During infection, *A. fumigatus* is predicted to use host carbon sources, such as acetate, present in body fluids and peripheral tissues, to sustain growth and promote colonisation and invasion. This work shows that *A. fumigatus* metabolises acetate via different pathways, a process that is dependent on the transcription factor FacB. Furthermore, the type and concentration of the extracellular available carbon source were determined to shape *A. fumigatus* virulence determinants such as secondary metabolite secretion and cell wall composition. Subsequently, interactions with immune cells are altered in a carbon source-specific manner. FacB is required for *A. fumigatus in vivo* virulence in both insect and mammalian models of invasive aspergillosis. This is the first report that characterises acetate utilisation in *A. fumigatus* and highlights the importance of available host-specific carbon sources in shaping virulence traits and potentially subsequent disease outcome.

## Introduction

*Aspergillus fumigatus* is a saprotrophic filamentous fungus and opportunistic pathogen of immunocompetent and immunocompromised hosts. Together with other opportunistic fungal pathogens, such as *Candida albicans* and *Cryptococcus neoformans*, globally they kill in excess of 1.5 million people a year(1). The severity of the diseases related to *A. fumigatus* depend on pre-existing infections as well as on the status of the host immune system (2). To successfully colonise and survive within the human host, *A. fumigatus* needs to acquire and metabolise nutrients. Essential nutrients include minerals such as iron, copper and zinc, which are required in small amounts; while carbon and nitrogen, the main energy sources for sustaining biosynthetic processes, must be obtained in large quantities(3). Iron, zinc and copper acquisition and metabolism have been studied in *A. fumigatus* in the context of virulence(4–6), whereas less is known about carbon source acquisition and metabolism in this fungus during infection. Studies have inferred that glucose, lactate and acetate are carbon sources available to fungi *in vivo* with their availability and concentration depending on the host niche (7, 8). Whereas glucose utilisation has been shown to be important for *A. fumigatus* disease progression (9), the utilisation of the physiologically relevant short chain fatty acids (SCFAs) lactate and acetate remain unexplored in this fungus. Indeed, acetate was detected in bronchoalveolar lavage (BAL) samples of healthy and immunosuppressed mice, suggesting the presence of this carbon source, independent of the underlying immune condition, at the *A. fumigatus* primary site of infection. Our work therefore aimed at characterising *A. fumigatus* acetate utilisation and its relevance for virulence.

In the human body, acetate is present in the blood plasma at concentrations ranging from 0.074 to 0.621 mM depending on the type of artery, diet and alcohol intake(10). Peripheral tissues can consume acetate from the blood stream and oxidise it(10). The main producer of plasma acetate is the gastrointestinal (GI) tract-resident microbiome, with the majority of GI-resident bacterial species being capable of producing acetate(10). Furthermore, acetate has immunoregulatory properties. Acetate is an agonist for the G-protein coupled receptors (GPCRs) FFA2, FFA3 and GPR109A, which are expressed in a number of immune cells, thus affecting the production of cytokines, the regulation of downstream anti-and pro-inflammatory responses, and recruitment of immune cells(11). Acetate is likely important during invasive fungal infections as it can regulate immunity at distal sites, including the lungs.

As mentioned above, small quantities of acetate were detected in healthy and immunosuppressed mice (8). The lungs are lined with a mucosa and contain a microbiome that has been shown to suffer alterations in the presence of disease(12). The lung microbiome, just like the gut microbiome, may contribute to the production and secretion of SCFAs(13). Studies investigating the production of SCFAs and other molecules by the lung microbiota are non-existent, probably due to the lungs having been thought of as sterile until a few years ago(12).

Our understanding of the utilization of potential food sources during infection mainly relies on *in vitro* transcriptional studies(14–16). Despite several studies having investigated *A. fumigatus* gene expression during *in vivo* infection of chemotherapeutic mice models of invasive aspergillosis, none of these studies have characterised the modulation of genes encoding components required for carbon source utilisation (17–20). The genome of *A. fumigatus* encodes the acetyl-CoA synthetases (ACS) FacA (Afu4g11080) and PcsA (Afu2g07780), with *facA* shown to be up-regulated in conidia exposed to neutrophils, and the corresponding protein induced by heat shock and repressed during hypoxic conditions(14, 21, 22). In *A. fumigatus*, the homologue of the *A. nidulans* transcription factor (TF)-encoding *facB* gene is up-regulated when conidia are exposed to neutrophils(14). Furthermore, *A. fumigatus* conidia which were exposed to human neutrophils from healthy or CGD (chronic granulomatous disease) donors, showed an up-regulation of genes encoding enzymes involved in the glyoxylate cycle, gluconeogenesis, peroxisome function and fatty acid degradation(14), suggesting an induction of metabolic pathways that are required for the utilisation of alternative, non-preferred carbon sources. Together, the aforementioned studies suggest that the utilisation of alternative carbon sources such as SCFAs is important for *A. fumigatus* infection.

Acetate utilisation has been investigated in detail in the model fungus *A. nidulans*. In *A. nidulans*, acetate was shown to be transported by the short-chain carboxylate transporters AcpA and AcpB, with the former being expressed in germinating conidia and young germlings, and the latter expressed in mycelia(23). The genome of *A. fumigatus* encodes one homologue of both *acpA* and *acpB* (Afu2g04080), which remains uncharacterised. Once internalised, acetate is converted by ACS to acetyl-CoA, which subsequently is transported into the mitochondria where it enters the TCA cycle for the synthesis of ATP molecules(24). Furthermore, acetate, in the acetyl-coA form, is oxidised via the glyoxylate cycle and required during gluconeogenesis(24). In *A. nidulans*, acetate utilisation is under the control of the TF FacB, which is transcriptionally induced in the presence of acetate(25). FacB controls the expression of the ACS FacA, of the carnitine acetyltransferases FacC, AcuH and AcuJ, of the succinate/fumarate antiporter AcuL and of the glyoxylate cycle malate synthase AcuE(25, 26). The genome of *A. fumigatus* encodes homologues of the *A. nidulans* components required for acetate metabolism, although they have not been investigated until now. This work characterised the utilisation of acetate in *A. fumigatus* and highlights the importance of the type of available extracellular carbon source in shaping fungal virulence determinants.

## Results

### Acetate is metabolised via the glyoxylate and TCA (tricarboxylic acid) cycles and a precursor for different metabolites

To investigate acetate metabolism in *A. fumigatus*, the metabolic fate of acetate was traced by incubating fungal mycelia with ^13^C_2_-labelled acetate. The *A. fumigatus* wild-type (WT) strain CEA17 was grown for 16 h in peptone rich-minimal medium (MM) before undergoing 4 h of carbon starvation in MM. Subsequently, ^13^C_2_-labelled acetate was added to the cultures for 5 and 15 min before mycelia were separated from the culture medium and immediately snap-frozen in liquid nitrogen. Following metabolite extraction, 1D ^1^H and 2D ^1^H-^13^C HSQC (heteronuclear single quantum coherence) NMR (nuclear magnetic resonance) spectra were recorded for each sample. The uptake of ^13^C_2_-acetate by *A. fumigatus* was observed through significant increases in carbon satellite peaks (reflecting ^1^H-^13^C coupling) on both sides of the central acetate singlet (δ 1.92 ppm) in ^1^H spectra of fungal cell extracts (Figure 1A, top). To identify metabolites that were directly derived from ^13^C_2_-labelled acetate and determine their fractional enrichment, ^1^H-^13^C HSQC spectra from 5 and 15 min samples were compared to spectra of control cells (4h starvation, no labelling). The extent of ^13^C-incorporation levels was obtained for each metabolite by dividing the 2D peak intensity in ^13^C-enriched samples by the peak intensity in the matched control sample (with 1.1% natural abundance in ^13^C levels). Results showed that labelled ^13^C from acetate was incorporated into the amino acids alanine at carbon-2 and carbon-3 (C2 and C3), aspartate at C2 and C3 and glutamate at C2, C3 and C4, the glyoxylate/TCA cycle intermediates citrate at C2, malate at C2 and C3 and succinate at C2 as well as in the glycolipid and glycoprotein compound N-acetylneuraminate at C11 (Figure 1A, bottom).

**Figure 1.**
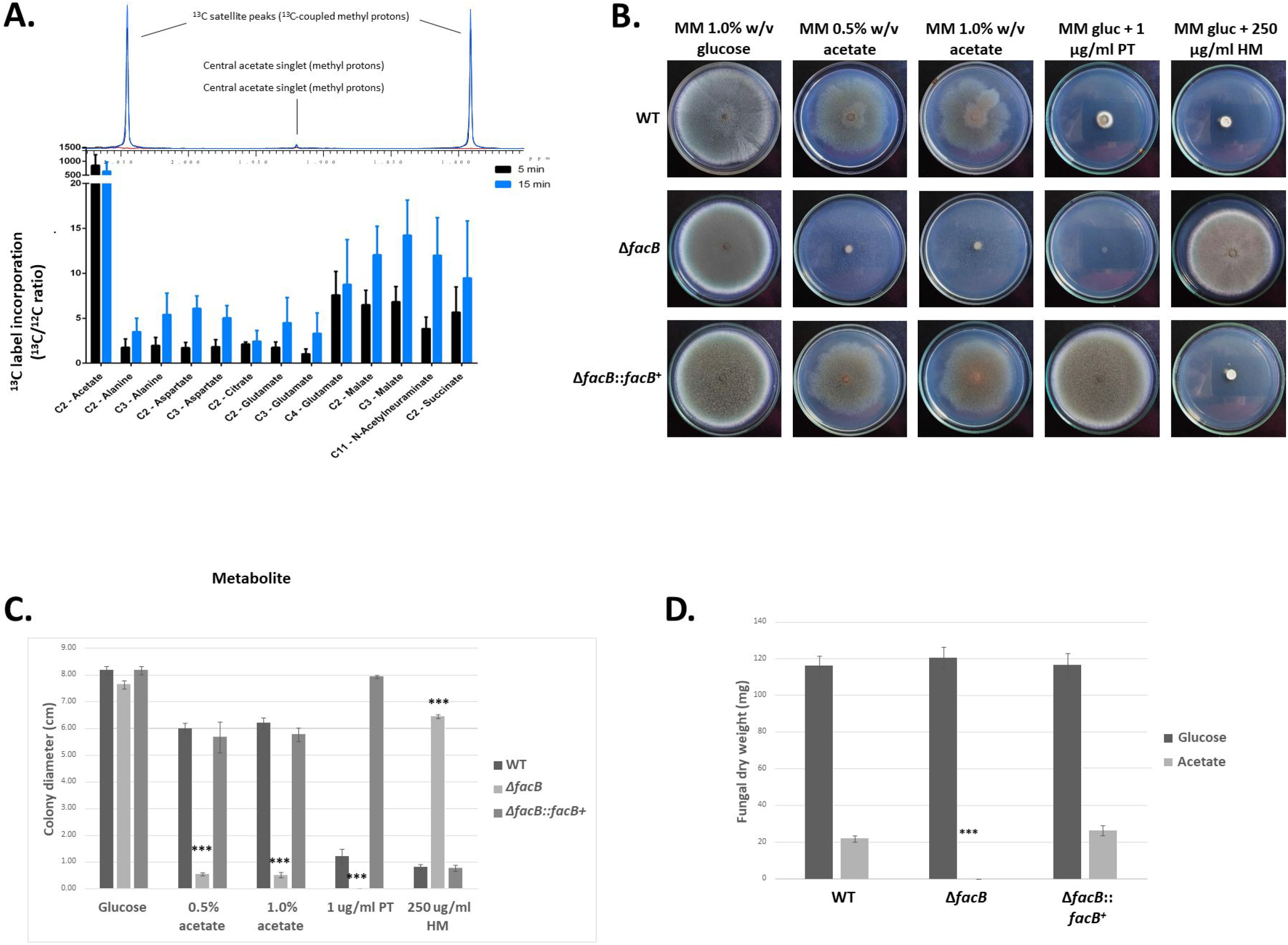
Acetate metabolism in *A. fumigatus*. **A**. One-and two-dimensional (1D, 2D) NMR (nuclear magnetic resonance) analysis of ^13^C_2_-labelled acetate incorporation and metabolism: (top) Expansion of 1D ^1^H NMR spectra of fungal cell extracts showing the increase in acetate carbon satellite peaks upon culture of *A. fumigatus* in ^13^C_2_-acetate-containing medium; (bottom) ^13^C/^12^C ratios for metabolites that incorporated ^13^C-derived from acetate, as determined through integration of ^1^H-^13^C HSQC (heteronuclear single quantum coherence) spectra recorded for extracts of fungal cells grown for 5 and 15 min in medium containing ^13^C_2_-acetate in comparison to non-labelled control cultures. Standard deviations represent the average of biological triplicates. **B-D**. The transcription factor FacB is essential for growth in the presence of acetate. Strains were grown for 5 days (B, C) or 3 days (D) in either solid (B, C) or liquid (D) minimal medium supplemented with 1% w/v glucose (gluc, B, D), 0.5% w/v (B) or 1% w/v (B, D) acetate before radial diameter (C) or fungal dryweight (D) was measured. To ensure homologous integration of *facB* in the complementation strain, strains were grown in the presence of pyrithiamine (PT, *facB* was re-introduced into the Δ*facB* strain using the PT-resistant marker gene) and hygromycin (HM, *facB* was deleted using the HM resistance marker gene) (B, C). Plate pictures (B) are representative for the average radial diameter shown in (C). Standard deviations represent the average of 3 biological replicates with \*\*\**p*-value < 0.0001 in a 2-way multiple comparisons ANOVA test, comparing the *facB* deletion strain to the WT strain.

These results indicate that acetate is taken up and metabolised via the glyoxylate and TCA cycles in *A. fumigatus*, which is in agreement with studies in *S. cerevisiae* (27) and *A. nidulans* (24) and in the line with the metabolism of two carbon compounds as the sole carbon source. Indeed, ^13^C was incorporated into the amino acid and Krebs cycle intermediate aspartate(28) as well as the amino acids alanine and glutamate. The TCA cycle intermediate oxaloacetate is converted into phosphor-enol-pyruvate (gluconeogenesis), which in turn is a precursor for alanine in a two-step process that involves glutamate. Furthermore, ^13^C was also enriched in other cellular compounds such as N-acetylneuraminate, which is a sialic acid that is present on cell surface acidic glycoconjugates, and which has been shown to contribute to the phagocytic properties of cells of opportunistic fungal pathogens such as *Cryptococcus neoformans* (29), *Candida albicans* (30) and *A. fumigatus* (31). Hence, this analysis showed that acetate is metabolised by *A. fumigatus* through multiple pathways.

### The transcription factor (TF) FacB is essential for *A. fumigatus* growth in the presence of acetate and ethanol as the sole carbon sources

To determine whether acetate utilisation is controlled by a TF in *A. fumigatus*, as was previously described for *A. nidulans* (25), a TF deletion library (32) was screened for reduced growth on plates containing MM supplemented with 0.5% (w/v) acetate (AMM) as the sole carbon source. Several strains were identified, and subsequent confirmation growth experiments, in both solid (radial growth) and liquid (dry weight) AMM, resulted in the selection of 5 strains that had reduced growth in acetate, but presented no growth defects in glucose-rich MM (GMM) (Supplementary Figure 1A-C at 10.6084/m9.figshare.14740482, Figure 1B-D). These strains were deleted for genes *acuK* (Afu2g05830), *acuM* (Afu2g12330), *facB* (Afu1g13510), *farA* (Afu4g03960) and *mtfA* (Afu6g02690) (Supplementary Figure 1 at 10.6084/m9.figshare.14740482, Figure 1B-D). AcuM, AcuK and MtfA have been characterised in *A. fumigatus* and have been shown to be important for alternative carbon source utilisation and virulence(33, 34). Furthermore, *farA* was shown to be important for fatty acid utilisation and was up-regulated in fungal cells exposed to human neutrophils(14). In contrast*, A. fumigatus* FacB, which is the homologue of *A. nidulans* FacB, remains uncharacterised. We therefore aimed at further deciphering the role of the TF FacB in *A. fumigatus* acetate utilisation and virulence. FacB was also essential for growth in medium with ethanol as the sole carbon source but not for growth in the presence of different fatty acids (Supplementary Figure 1D at 10.6084/m9.figshare.14740482). Ethanol and acetate are two carbon compounds that require identical metabolic pathways with ethanol being converted to acetate via the metabolic intermediate acetaldehyde(35). Re-introduction of *A. fumigatus facB* in the Δ*facB* background strain at the *facB* locus through homologous recombination restored growth in acetate (Figure 1B-D, Supplementary Figure 1D at 10.6084/m9.figshare.14740482), confirming that the FacB-encoding gene is essential for *A. fumigatus* growth on two carbon compounds.

### FacB controls acetate utilisation through regulating genes encoding enzymes required for acetate metabolism

To gain further insight into *A. fumigatus* acetate metabolism and to describe a role of FacB in the control of acetate utilisation, the transcriptional response of the wild-type (WT) and Δ*facB* strains was assessed by RNA-sequencing (RNA-seq), when grown for 24 h in fructose-rich (control) MM and after transfer for 0.5 h (short incubation) or 6 h (long incubation) to MM supplemented with 0.1% w/v (low concentration) or 1% w/v (high concentration) acetate. We chose different concentrations of acetate and time points in order to decipher the transcriptional response in the presence of abundant and limiting carbon source concentrations after short and prolonged exposure. The number of significantly differentially expressed genes (DEGs) were defined as having a −1 < log2FC (fold change) < 1 and an adjusted *p*-value < 0.05 (Supplementary File 1 at 10.6084/m9.figshare.14740482, Table 1). Two comparisons were carried out: i) gene expression in the presence of the four different acetate conditions against gene expression in the control (fructose) condition in the WT strain; and ii) gene expression in the WT strain against gene expression in the Δ*facB* strain in the presence of the different acetate conditions (Table 1, 8 comparisons in total). Gene ontology (GO) and Functional Categorisation (FunCat) analyses could not be performed for many of the comparisons shown in Table 1, probably due to a low number of DEGs in some conditions (Table 1). DEGs were therefore manually inspected and divided into the following categories: a) amino acid, protein and nitrogen (urea, nitrate, ammonium) metabolism (degradation and biosynthesis); b) carbohydrate and lipid metabolism, including genes encoding enzymes required for lipid, fatty acid and acetate degradation, CAZymes (carbohydrate active enzymes) and the metabolism of other sugars; c) cell signalling (protein kinases, phosphatases, regulators of G-protein signalling and G-protein coupled receptors – GPCRs); d) cell membrane and cell wall (ergosterol, chitin and glucan biosynthesis/degradation); e) miscellaneous (genes encoding enzymes with diverse functions that do not fit into the other categories); f) oxidation/reduction and respiration (oxidoreductases, monooxygenases and respiratory chain enzymes); g) secondary metabolism; h) transcription factors; i) transporters (sugars, amino acids, ammonium, nitrate, ions, metals and multidrug); j) unknown (gene encoding proteins with unknown/uncharacterised functions) and k) putative virulence factors (proteases and proteins important for adhesion and interaction with the extracellular environment) (Supplementary Files 2 and 3 at 10.6084/m9.figshare.14740482; Supplementary Figure 2 at 10.6084/m9.figshare.14740482). In the WT strain, this categorisation was carried out for all DEGs with a −3 < log2FC < 3 in order to identify genes with the highest differential expression pattern. For comparisons between the WT and Δ*facB* strains, categorisation was carried out for all DEGs with a −1.5 < log2FC < 1.5 in order to include as many DEGs as possible.

**Table 1.**
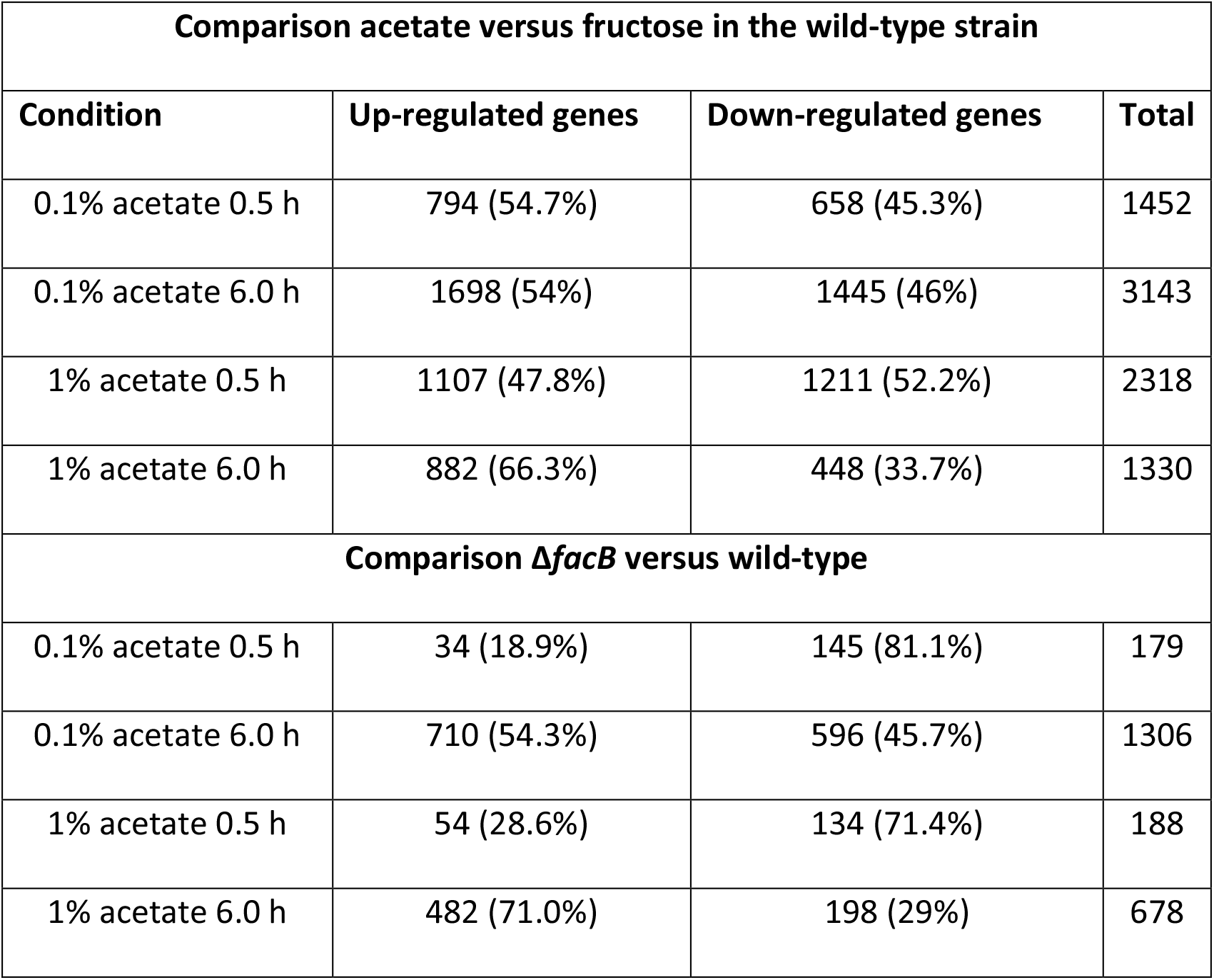
Number of differentially expressed genes (DEGs, −1 < log2FC < 1) identified by RNA-sequencing in the wild-type and Δ*facB* strains when grown for 0.5 h or 6 h in minimal medium supplemented with 0.1 or 1.0 % w/v acetate.

The majority of DEGs (34 – 46%) encoded proteins of unknown function, whereas genes encoding enzymes required for carbohydrate and carbon compound (CC) metabolism, oxidation/reduction and respiration, secondary metabolism and transporters constituted 38 – 48% of all DEGs (Supplementary Figure 2 at 10.6084/m9.figshare.14740482) suggesting the presence of acetate influences the regulation of these processes. No particular enrichment for any of the aforementioned categories was found for the here studied conditions (Supplementary Figure 2 at 10.6084/m9.figshare.14740482). There were differences though in the type of secondary metabolites (SMs), transporters as well as respiratory and carbon source metabolism encoded by the DEGs (Supplementary Figure 2 at 10.6084/m9.figshare.14740482).

To further unravel the role of FacB in acetate utilisation, we focused on DEGs that encode enzymes important for acetate metabolism. In the wild-type strain, genes encoding the ACS FacA (but not the ACS PcsA), a carnitine acetyl transferase (Afu1g12340), a mitochondrial carnitine:acyl carnitine carrier (Afu6g14100), the isocitrate lyase (ICL) AcuD (*acuD*, glyoxylate cycle) and the malate synthase AcuE (*acuE*, glyoxylate cycle) were highly expressed in all acetate conditions; in contrast, these genes were repressed in the Δ*facB* strain (Figure 2A). The exception was in the presence of 6 h in 0.1% w/v acetate, where these genes were not expressed in the WT strain but they were induced in the *facB* deletion strain (Figure 2A). This is likely due to carbon starvation, which would occur in these conditions. Assessment of the expression of *facA*, Afu1g12340, Afu6g14100 and *acuD* by qRT (real-time reverse transcriptase)-PCR in the same conditions confirmed the RNA-seq data (Figure 2B, 2C).

**Figure 2.**
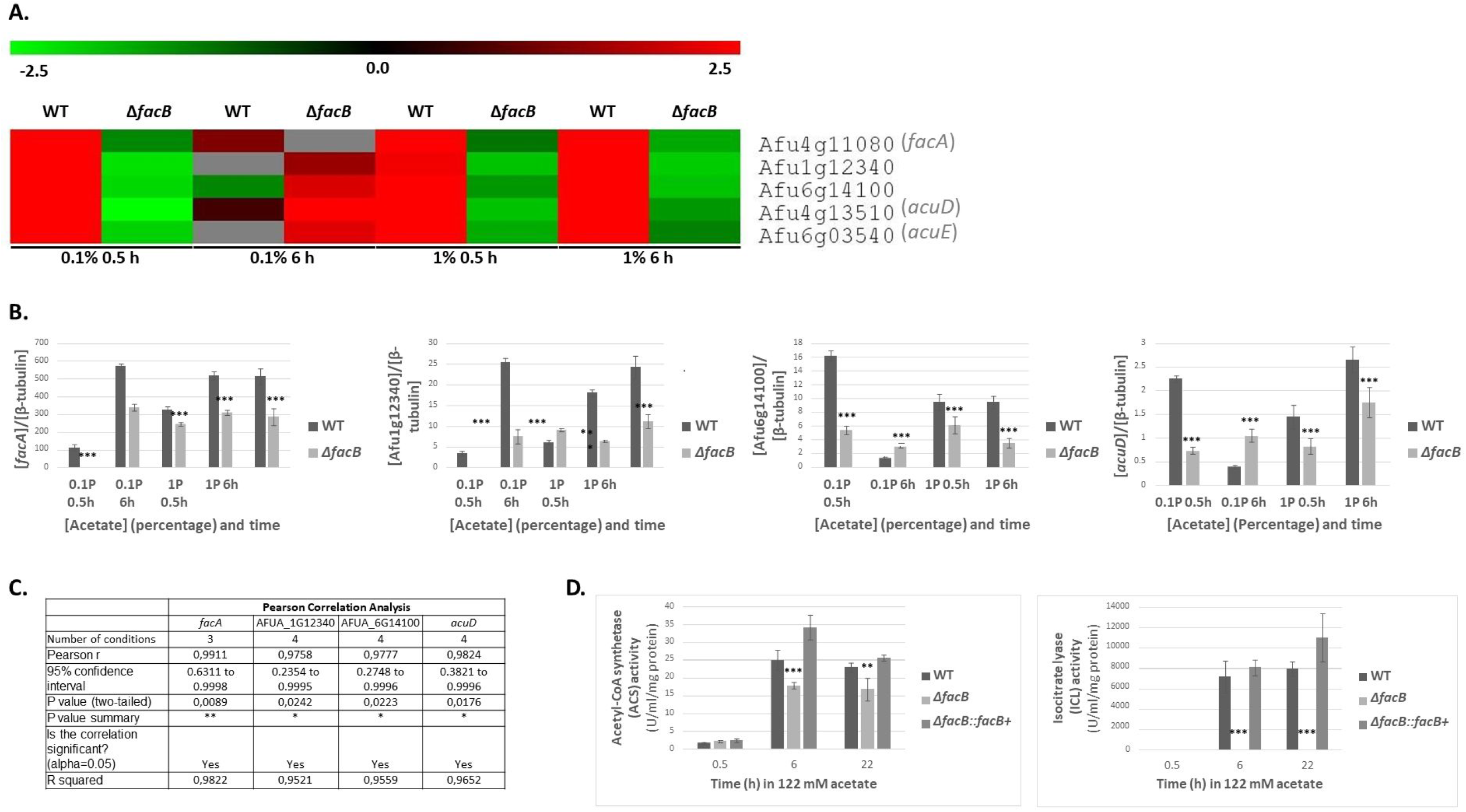
FacB regulates acetate metabolism. **A**. Heat map depicting log2FC (fold changes) from the RNA-sequencing data of genes encoding enzymes required for acetate metabolism in the wild-type (WT) and Δ*facB* strains in the presence of 0.1% w/v or 1% w/v acetate after 0.5 h and 6 h. The log2FC for the WT strain is based on the comparison of gene expression between the WT strain grown in fructose-rich medium and after transfer to acetate-rich medium; whereas log2FC for the Δ*facB* strain is from the comparison between the WT and *facB* deletion strain for each acetate condition. **B**. Validation of RNA-sequencing data by qRT-PCR shows that FacB is required for the transcriptional expression of genes encoding enzymes involved in acetate metabolism. Strains were first grown in minimal medium (MM) supplemented with fructose before mycelia were transferred to acetate-containing MM, RNA was extracted and reverse-transcribed to cDNA and qRT-PCR was run on genes *facA*, Afu1g12340, Afu6g14100 and *acuD*. Gene expressions were normalised by β-tubulin. **C.** Results of Pearson Correlation Analysis between the RNA-sequencing and qRT-PCR datasets for 4 genes encoding enzymes involved in acetate metabolism. Gene fold-change values were used for the analysis, which was carried out in Prism Graphpad (\**p*-value < 0.05, \*\**p*-value < 0.005). **D**. FacB is required for acetyl-CoA synthetase (ACS) and isocitrate lyase (ICL) activities. Strains were grown in fructose-rich MM for 24 h, before mycelia were transferred to MM containing 1% w/v acetate for 0.5 h, 6 h and 22 h and total cellular proteins were extracted and enzyme activities were measured. Standard deviations represent the average of 3 biological replicates with \*\**p*-value < 0.001, \*\*\**p*-value < 0.0001 in a 2-way multiple comparisons ANOVA test when comparing the *facB* deletion strain to the WT strain.

To further confirm the transcriptional data, we assayed the activities of ACS and ICL (isocitrate lyase) in the WT, Δ*facB* and Δ*facB*::*facB*^+^ strains when grown in the presence of 1% w/v acetate for 0.5 h, 6 h and 22 h. In agreement with the RNA-seq data, ACS and ICL activities were induced in the presence of acetate in the WT and Δ*facB*::*facB*^+^ strains. No significant difference in ACS and ICL activities were observed between the WT and Δ*facB*::*facB*^+^ strains, whereas these enzyme activities were significantly reduced in the Δ*facB* strain in all tested conditions (Figure 2D). ICL activity was completely dependent on FacB with the loss of *facB* resulting in no enzyme activity (Figure 2D). In contrast, ACS activity was reduced ∼20-30% in the Δ*facB* strain when compared to the WT and Δ*facB*::*facB*^+^ strains (Figure 2D). The observed ACS activity is likely due to the activity of the second *A. fumigatus* ACS PcsA. Our RNA-seq data shows that *pcsA* is not under the regulatory control of FacB in the here tested conditions, whereas the expression of the single ICL-encoding gene *acuD*, is regulated by FacB (Figure 2A). These results suggest that FacB controls acetate utilisation through regulating genes encoding enzymes required for acetate metabolism.

### Acetate metabolism is subject to carbon catabolite repression (CCR)

In *A. fumigatus*, CCR is a cellular process which directs primary metabolism to the utilisation of preferred carbon sources (glucose) and results in the repression of genes required for the utilisation of alternative carbon sources (acetate)(36). The opportunistic yeast pathogen *C. albicans* is able to simultaneously use glucose and lactate, due to the loss of an ubiquitination site on ICL(37). This increased metabolic flexibility plays a role in the adaptation of *C. albicans* to the host environment with the addition of an ubiquitination site to *C. albicans* ICL resulting in decreased resistance to phagocytosis by macrophages, decreased fungal burden in the GI tract, and decreased dissemination to the kidneys(38). To determine whether *A. fumigatus* is able to use glucose and acetate simultaneously, transcriptional and enzymatic studies were performed in the WT and Δ*facB* strains in the presence of equimolar concentrations of glucose and acetate. We also included a strain deleted for the TF CreA, which is a transcriptional regulator of CCR (36). Strains were grown in the presence of 12.2 mM (0.1% w/v) and 122 mM (1% w/v) acetate without or with equimolar concentrations of glucose for 0.5 h, before expression of genes *facB*, *facA*, Afu1g12340, Afu6g14100 and *acuD* were determined by qRT-PCR (Figure 3A-B). The presence of the different concentrations of glucose caused a significant down-regulation of all genes, except for *facB* in the presence of 122 mM acetate and glucose (Figures 3A-B). It is possible that the expression of *facB* is dependent on the concentration of the externally available carbon source. In *Aspergillus spp*., high and low affinity carbon source transporters are expressed depending on the concentration of the extracellular carbon source (39). A similar scenario can be envisaged for transcription factors (TFs), especially as they respond to external stimuli, with some TFs being highly induced under nutrient limiting conditions and repressed in nutrient sufficient conditions (20). Alternatively, *Aspergillus* transcriptions factors are not always regulated at the transcriptional level as previously shown (40). These results suggest that *A. fumigatus* acetate metabolism is subject to CCR as has been described in *A. nidulans*(25, 26).

**Figure 3.**
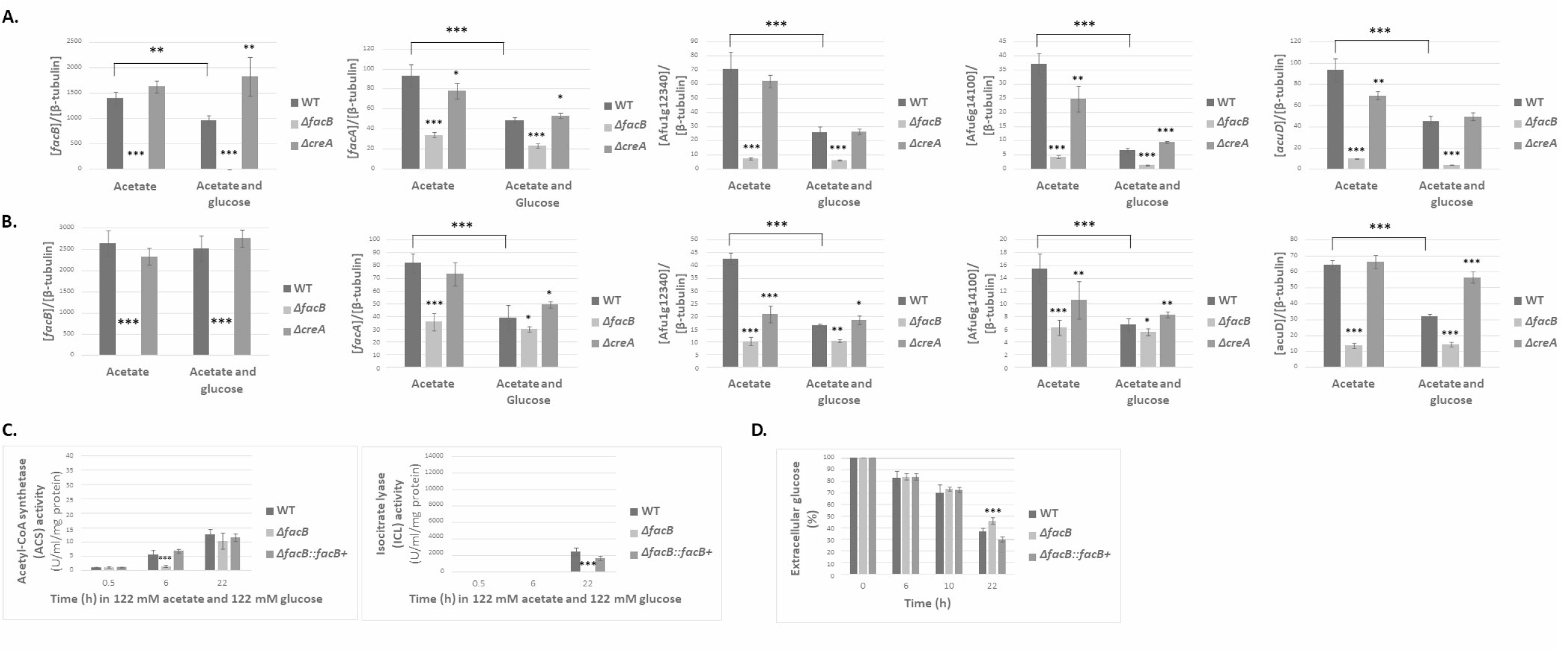
Acetate utilisation is subject to carbon catabolite repression (CCR). **A**. – **B**. Expression of genes *facA*, Afu1g12340, Afu6g14100 and *acuD*, as determined by qRT-PCR, in strains grown for 24 h in minimal medium (MM) supplemented with fructose and then transferred for 0.5 h to MM supplemented with either acetate or acetate and glucose. Graphs in panel A. show results from growth in 12.2 mM for each carbon source whereas graphs in panel B. show results from growth in 122 mM for each carbon source. **C**. Activities of acetyl-CoA synthetase (ACS) and isocitrate lyase (ICL) in strains incubated for 0.5 h, 6 h and 22 h in MM supplemented with 122 mM acetate and 122 mM glucose. Strains were first grown for 24 h in fructose MM before mycelia were transferred to acetate and glucose-containing MM. **D**. Percentage of residual glucose in supernatants of strains grown for 24 h in fructose-rich MM and after transfer to glucose MM for a total time period of 22 h. Standard deviations represent the average of 3 biological replicates with \**p*-value < 0.01, \*\**p*-value < 0.001, \*\*\**p*-value < 0.0001 in a 2-way multiple comparisons ANOVA test when comparing the *facB* deletion strain to the WT strain or when comparing the WT strain in two conditions (indicated by a line).

In the presence of 12.2 mM acetate, deletion of *creA* caused a significant down-regulation of *facA*, Afu6g14100 and *acuD*; whereas in the simultaneous presence of 12.2 mM acetate and glucose, the absence of *creA* significantly increased *facB* and Afu6g14100 gene expression (Figure 3A). In the presence of 122 mM acetate, deletion of *creA* significantly reduced Afu1g12340 and Afu6g14100 gene expression, whereas upon in the presence of glucose, the expression of all genes, except for *facB*, was increased, although not to WT levels (Figure 3B). The exception was the expression of *acuD* in the Δ*creA* strain in the presence of 122 mM acetate and glucose, which was similar to the expression levels of *acuD* in the WT strain in the presence of 122 mM acetate (Figure 3B). These results suggest that: i) CreA may be involved in the control of genes required for alternative carbon source utilisation and that ii) acetate metabolism (with the exception of *acuD*) is partially dependent on CreA-mediated CCR in a concentration-dependent manner.

Next ACS and ICL activities were measured in the presence of 122 mM acetate and glucose after 0.5 h, 6 h and 22 h. Enzyme activities were lower in the presence of acetate and glucose (Figure 3C) than in the presence of acetate only (Figure 2C), supporting the observed transcriptional repression of the corresponding genes in the presence of glucose. Basal ACS activity was detected in all conditions (likely due to the presence of the FacB-dependent ACS FacB and the FacB-independent ACS PcsA) whereas ICL activity was not detected at 0.5 h and 6 h. This is in agreement with the transcriptional data, suggesting that ICL activity is completely dependent on FacB for induction (Figures 2B-C, 3B-C) and CreA for repression (Figure 3B). After 22 h incubation in both carbon sources, enzyme activities increased, which may be due to low glucose concentrations (∼ 30%) in the culture medium (Figure 3D), making acetate the predominant available carbon source. Furthermore, significantly more extracellular glucose was present in supernatants of the Δ*facB* strain (Figure 3D), suggesting that FacB may also be involved in the utilisation of other carbon sources. Together, these results suggest that genes and enzymes involved in acetate utilisation are subject to CCR and that their regulation is partially controlled by CreA.

### The extracellular carbon source influences the levels of secreted secondary metabolites (SMs)

The secretion of SMs has been shown to be essential for *A. fumigatus* proliferation within the natural environment and mammalian host, for evasion and modulation of the host immune system and for virulence(41). Our RNA-seq data shows that many DEGs are part of the fumagillin, pseurotin A, pyomelanin and gliotoxin SM biosynthetic gene clusters (BGCs) (Supplementary Figure 2 at 10.6084/m9.figshare.14740482, Figure 4A-B). SM BGCs are mainly expressed in the WT strain in the presence of 1% w/v acetate or after 6 h incubation in MM supplemented with 0.1% w/v acetate (Figure 4A-B). In contrast, these DEGs have reduced expression or are repressed in the in the Δ*facB* strain (Figure 4A-B). Furthermore, the expression profiles of these genes were often reversed between the WT and Δ*facB* strains in these conditions, suggesting that the metabolic pathways regulated by FacB are important for SM gene expression.

**Figure 4.**
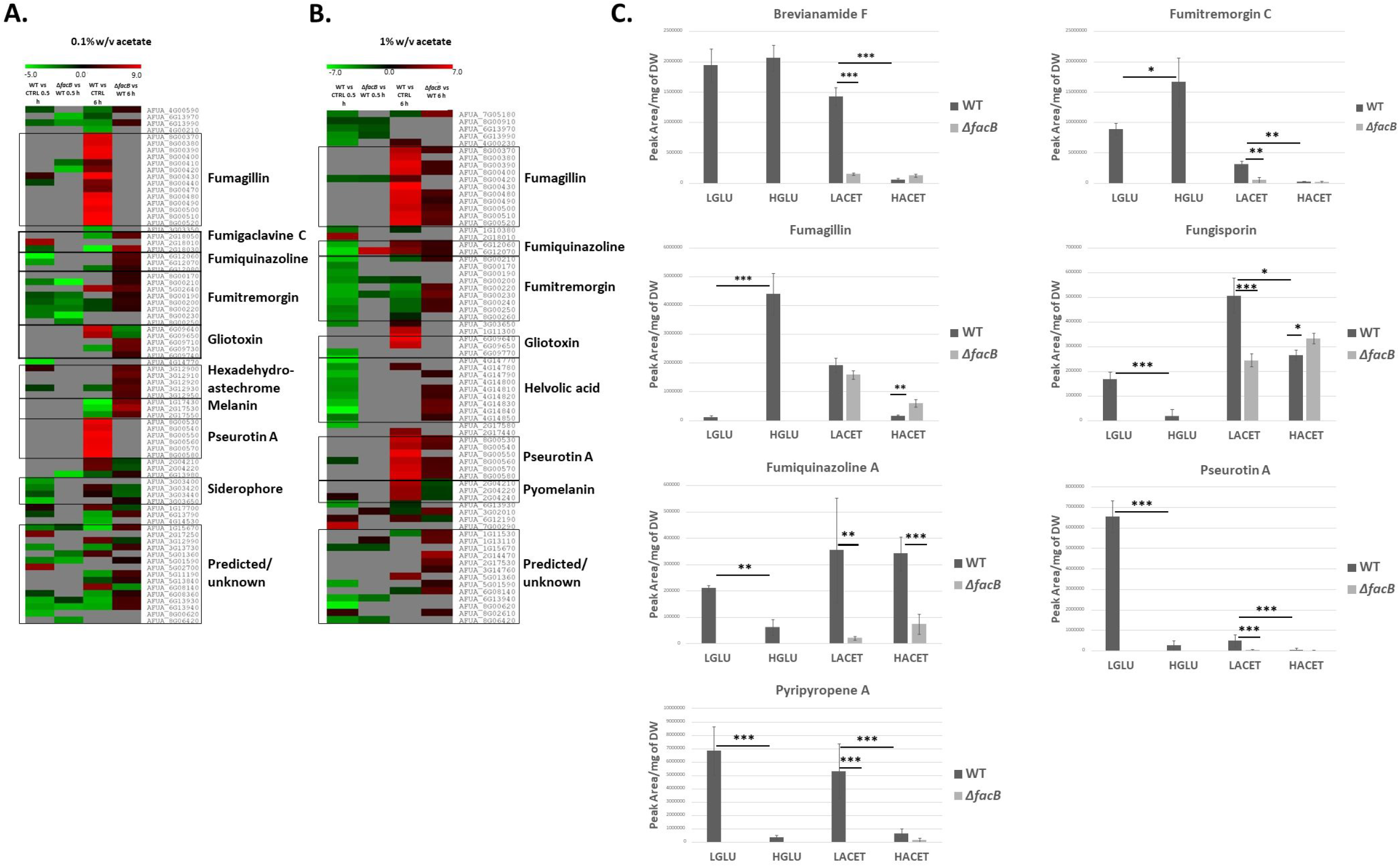
The extracellular carbon source affects the levels of secreted secondary metabolites (SMs). **A, B**. Heat map of the log2 fold-change (FC), as determined by RNA seq, of genes predicted to encode enzymes required for SM biosynthesis in the wild-type (WT) and Δ*facB* strains when grown for 0.5 h and 6 h in the presence of 0.1% w/v or 1% w/v acetate or when comparing gene expression in the WT strain in the presence of different acetate concentrations and in the presence of fructose (control, CTRL condition). In grey, are genes that did not show a significant FC. **C**. Quantities of identified SMs, as determined by high performance liquid chromatography (HPLC), in the WT and Δ*facB* strains when grown for 24 h in minimal medium supplemented with 0.1% w/v (LGLU = low glucose; LACET = low acetate) or 1% w/v glucose (HGLU = high glucose; HACET = high acetate) or acetate. SM quantities were normalised by fungal dry weight (DW). Standard deviations represent the average of 4 biological replicates with \**p*-value < 0.01, \*\**p*-value < 0.001, \*\*\**p*-value < 0.0001 in a 2-way multiple comparisons ANOVA test.

To determine whether SMs are secreted specifically in the presence of acetate and dependent on FacB, high performance liquid chromatography (HPLC) was performed on culture supernatants from the WT and Δ*facB* strains grown for 24 h in fructose-rich MM and after transfer to MM supplemented with 0.1% w/v and 1% w/v acetate for 24 h. This pre-growth in fructose ensured that the starting biomass was similar for all samples. In addition, SM profiles were examined for the WT strain when grown in the same conditions, with the exception that acetate was replaced with glucose as the main carbon source. After 24 h, a total of 18 SMs including the previously characterised(42, 43) fumiquinazolines A and D, fumitremorgin C, pyripyropene A, pseurotins A and F2, fungisporin and brevianamide F, were identified in culture supernatants from strains grown in all conditions (Supplementary Table 2 at 10.6084/m9.figshare.14740482). We did not detect gliotoxin or pyomelanin in culture supernatants.

Subsequently, the concentrations of characterised SMs were quantified to determine whether the extracellular available carbon source influences the levels of secreted SMs. In the WT strain, concentrations of all these SMs, with the exception of fumiquinazoline and fungisporin, were significantly higher in the presence of 1% w/v glucose than in the presence of 1% w/v acetate (Figure 4C). Similarly, in the presence of 0.1% w/v glucose, secreted levels of brevianamide F, fumiquinazoline, fumitremorgin C and pseurotin A were significantly higher when compared to concentrations in the presence of 0.1% w/v acetate; whereas levels of fumagillin and fungisporin A were significantly higher in the presence of 0.1% w/v acetate than in the presence of 0.1% glucose (Figure 4C). Furthermore, differences in levels of secreted SMs were also seen between the two different concentrations of the same carbon source (Figure 4C). In addition, deletion of *facB* resulted in a significant decrease in concentrations of secreted SMs in the presence of different concentrations of extracellular acetate with the exception of fumagillin (Figure 4C).

Together, these results suggest that the concentration and type of available extracellular carbon source affects the levels of secreted SMs.

### The composition of the *A. fumigatus* cell wall is carbon source dependent

In *A. fumigatus*, the composition of the culture medium influenced cell wall composition, thus modulating their sensitivity to antifungal agents(44). In addition, primary carbon metabolism was shown to influence cell wall content and/or organisation(45). To investigate whether glucose and acetate, respectively representing preferred and alternative carbon sources would also influence cell wall composition, we determined the quantities of cell wall polysaccharides in the *A. fumigatus* WT strain when grown in the presence of each of these carbon sources.

Cell wall alkali-insoluble (AI) and alkali-soluble (AS) fractions were prepared of WT mycelia grown for 24 h in MM supplemented with 1% w/v glucose or acetate and analysed by gas-liquid chromatography (Supplementary Figure 3A at 10.6084/m9.figshare.14740482). Results show that there is a significant increase in the percentage of the cell wall AI fraction in the presence of acetate due to increased concentrations of glucose (β-1,3-glucan) and glucosamine (chitin) (Figures 5A and 5B; Supplementary Figure 3A at 10.6084/m9.figshare.14740482). In contrast, the percentage of cell wall AS fraction was significantly reduced in the presence of acetate predominantly due to decreased levels of glucose (α-1,3-glucan), although significantly decreased levels of mannose and galactose were also observed (Figure 5A,B; Supplementary Figure 3A at 10.6084/m9.figshare.14740482). These results suggest that the type of extracellular carbon source significantly influences *A. fumigatus* cell wall composition and organisation.

**Figure 5.**
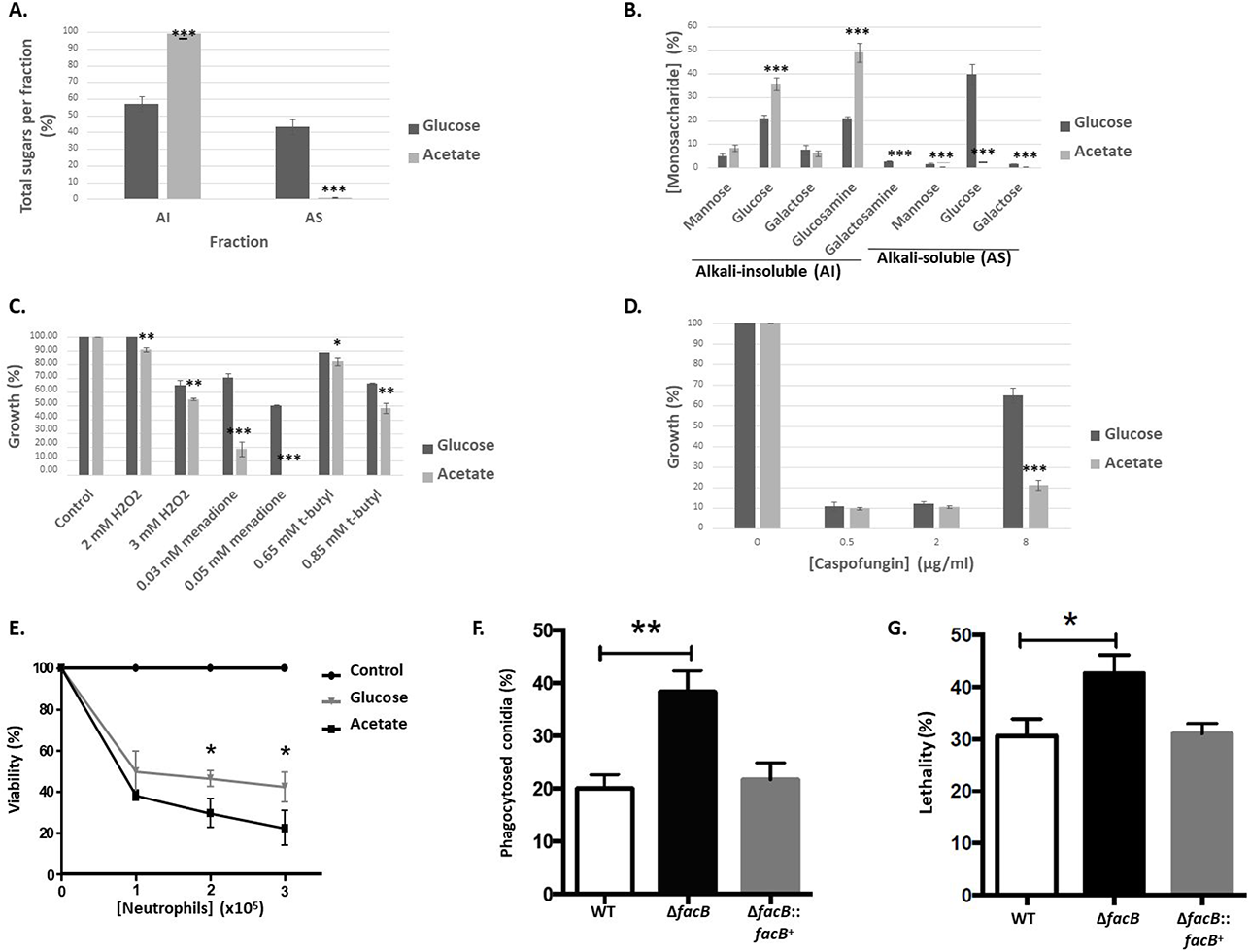
Acetate utilisation impacts cell wall polysaccharide content, oxidative stress, caspofungin and immune cell resistance in *A. fumigatus*. **A, B.** Percentage of total (A.) and individual (B.) sugars identified in the alkali-insoluble (AI) and alkali-soluble (AS) fractions by gas liquid chromatography of the WT strain when grown for 24 h in MM supplemented with glucose and acetate. Standard deviations represent the average of 4 biological replicates and \*\*\**p*-value < 0.0001 in a 2-way multiple comparisons ANOVA test when comparing the acetate condition to the glucose condition. **C**. The wild-type (WT) strain was grown from 10^5^ spores for 5 days on glucose (GMM) or acetate minimal medium (AMM) supplemented with different concentrations of oxidative stress-inducing compounds. Colony diameters were measured and normalised by the control condition and expressed as percentage of growth in comparison to the control condition. **D**. As described in C., with the exception that GMM or AMM was supplemented with increasing concentrations of caspofungin. Standard deviations represent the average of 3 biological replicates and \*\**p*-value < 0.001, \*\*\**p*-value < 0.0001 in a 2-way multiple comparisons ANOVA test comparing the acetate condition to the glucose condition. **E.** The WT strain was pre-grown for 8 h or 13 h in GMM or AMM respectively, before hyphae were incubated with different concentrations of human neutrophils for 1 h. Subsequently, cells were lysed and hyphal viability was assessed via an MTT assay and calculated. Standard deviations represent the average of 3 biological replicates and \**p*-value < 0.05 in a one-tailed t-test, comparing the acetate condition to the glucose condition. **F, G.** Murine bone marrow-derived macrophage (BMDM) phagocytosis (F.) and killing (G.) of *A. fumigatus* conidia. BMDMs were incubated with fungal conidia before they were stained with calcofluor white and the percentage of phagocytosed conidia was assessed by microscopy and calculated. To assess fungal viability, conidia-macrophage mixtures were lysed, diluted and inoculated on plates containing complete medium before colony forming units (CFU) were assessed and percentage of viability calculated. Standard deviations represent the average of 3 biological replicates and \**p*-value < 0.05, \*\**p*-value < 0.005 in a paired t-test.

### Oxidative stress and antifungal drug tolerance are carbon source-dependent

The *A. fumigatus* cell wall has been shown to play a major role during infection as it represents the main line of defence for the fungus and is responsible for interacting with and modulating host immune cells as well as for oxidative stress and antifungal drug resistance(46). The aforementioned results show that the type of carbon source has an effect on cell wall polysaccharide concentrations. Subsequently, oxidative stress and antifungal drug resistance were determined in the *A. fumigatus* WT when grown in the presence of glucose or acetate.

First, the WT strain was grown in the presence of GMM or AMM supplemented with the oxidative stress-inducing compounds hydrogen peroxide (H_2_O_2_), menadione and *t*-butyl hydroperoxide, before colony diameters were measured and the percentage of growth was normalised by the growth in the control, drug-free condition for each carbon source. Growth was significantly reduced in the presence of AMM supplemented with the oxidative stress-inducing compounds when compared to growth in the presence of GMM supplemented with the oxidative stress-inducing compounds (Figure 5C, Supplementary Figure 3B at 10.6084/m9.figshare.14740482). These results suggest that the presence of acetate increases sensitivity to oxidative stress in *A. fumigatus*.

Next, we determined resistance to antifungal drugs, including different azoles, amphotericin B and caspofungin, when the WT strain was grown in the presence of glucose and acetate. We performed MIC (minimal inhibitory concentration) assays of amphotericin B, voriconazole, itraconazole and posaconazole when *A. fumigatus* was grown in RPMI medium (standard reference medium used for MIC assays), GMM and AMM for 72 h. *A. fumigatus* grown in the presence of AMM, was slightly more susceptible to amphotericin B than when compared to growth in the presence of RPMI and GMM (Table 2). No difference in susceptibility was observed for the different here tested azoles when compared to RPMI although reduced growth in the presence of these azoles was observed when comparing MIC between GMM and AMM (Table 2). The WT strain was also grown in GMM or AMM supplemented with increasing concentrations of the echinocandin and second line therapy drug caspofungin (47). In the presence of 0.5 and 2 μg/ml caspofungin, growth was severely inhibited and did not differ between both carbon sources (Figure 5D, Supplementary Figure 3C at 10.6084/m9.figshare.14740482). At 8 μg/ml caspofungin, the WT strain had increased growth in the presence of glucose and acetate, due to the caspofungin paradoxical effect (increased fungal growth in the presence of higher caspofungin concentrations (48)). In the presence of acetate, the WT was less able to recover growth when compared to growth on GMM (Figure 5D, Supplementary Figure 3C at 10.6084/m9.figshare.14740482).

**Table 2.** Minimal inhibitory concentrations (MIC) of different antifungal drugs (μg/ml) on the *A. fumigatus* WT strain when grown in RPMI, glucose (GMM) and acetate minimal medium (AMM). Shown is the average and standard deviations of three independent repeats with * *p* < 0.05 in a two-way ANOVA test comparing AMM to RPMI and GMM.

| Medium | Amphotericin B | Voriconazole | Itraconazole | Posaconazole |
| --- | --- | --- | --- | --- |
| RPMI | $3.33 \pm 1.15$ | $0.25 \pm 0.00$ | $0.42 \pm 0.14$ | $0.67 \pm 0.29$ |
| GMM | $3.33 \pm 1.15$ | $0.33 \pm 0.14$ | $0.50 \pm 0.00$ | $1.00 \pm 0.00$ |
| AMM | $1.17^* \pm 0.76$ | $0.21 \pm 0.07$ | $0.33 \pm 0.14$ | $0.67 \pm 0.29$ |

These results suggest that oxidative stress and antifungal drug resistance change depending on the extracellular, available carbon source.

### Acetate-grown hyphae are more susceptible to human neutrophil-mediated killing than hyphae grown in the presence of glucose

The aforementioned results indicate that growth in the presence of acetate influences virulence determinants such as SM production, cell wall composition, oxidative stress and antifungal drug resistance when compared to growth in the presence of energetically more favourable carbon sources such as glucose. To determine the role of carbon source-mediated growth for resistance against human neutrophils, we assayed the viability of hyphae, pre-grown in MM supplemented with either glucose or acetate as the sole carbon source. To ensure that a similar number of conidia had germinated prior to incubation with neutrophils, microscopy was performed and the number of germinated conidia was counted. After incubation for 8 h in GMM and 13 h in AMM, ∼ 90% of conidia had germinated in both conditions (Supplementary Figure 3D at 10.6084/m9.figshare.14740482) and they were visually inspected to be similar in length (data not shown). Human neutrophils used at different multiplicity of infection (MOI), killed significantly more (60 - 80%) *A. fumigatus* hyphae grown in AMM than when compared to hyphae (50 - 60%) pre-grown in GMM (Figure 5E). These results indicate that hyphae grown in acetate-rich medium are more susceptible to human neutrophil-mediated killing than hyphae grown in the presence of glucose.

### FacB is crucial for virulence in insect and murine models of disseminated and invasive pulmonary aspergillosis (IPA)

Lastly, we assessed the virulence of the Δ*facB* strain in *vitro* and *in vivo*. First, the capacity of murine bone marrow-derived macrophages (BMDM) to phagocytose and kill WT, Δ*facB* and Δ*facB::facB*^+^ conidia was determined. The Δ*facB* strain was significantly more susceptible to BMDM phagocytosis (Fig. 5F) and a significantly higher amount of Δ*facB* conidia were killed in comparison to the WT and Δ*facB::facB*^+^ strains (Fig. 5G).

Next, virulence of the WT, Δ*facB* and Δ*facB::facB*^+^ strains was determined in the wax moth *Galleria mellonella* and in a neutropenic murine model of IPA. We used different animal models as virulence *A. fumigatus* was shown to be dependent on the status of the host immune system(49). In *G. mellonella* (Figure 6A) and in chemotherapeutic mice (Figure 6B), the Δ*facB* strain was hypovirulent when compared to the WT and Δ*facB::facB*^+^ strains. In the insect model, the WT and Δ*facB::facB*^+^ strains killed all larvae after 8 days, whereas 80% of larvae infected with the Δ*facB* strain survived after 10 days (Figure 6A). Similarly, the WT and Δ*facB::facB*^+^ strains killed all mice after 4 days, whereas 10% of mice infected with the Δ*facB* strain survived 6 days post-infection (p.i) (Figure 6B). In agreement, fungal burden was significantly reduced for the Δ*facB* strain after 3 (Figure 6C) and 7 (Figure 6D) days p.i. in murine lungs when compared to the WT and Δ*facB::facB*^+^ strains. In addition, histopathology analyses of murine lungs after 3 p.i., showed significantly reduced inflammation (Figure 6E, F) and growth in the lungs (Figure 6F) for the Δ*facB* strain after 3 days p.i. when compared to the WT and Δ*facB::facB*^+^ strains. Together these results suggest that FacB is important for *A. fumigatus* virulence in insect and mammalian hosts.

**Figure 6.**
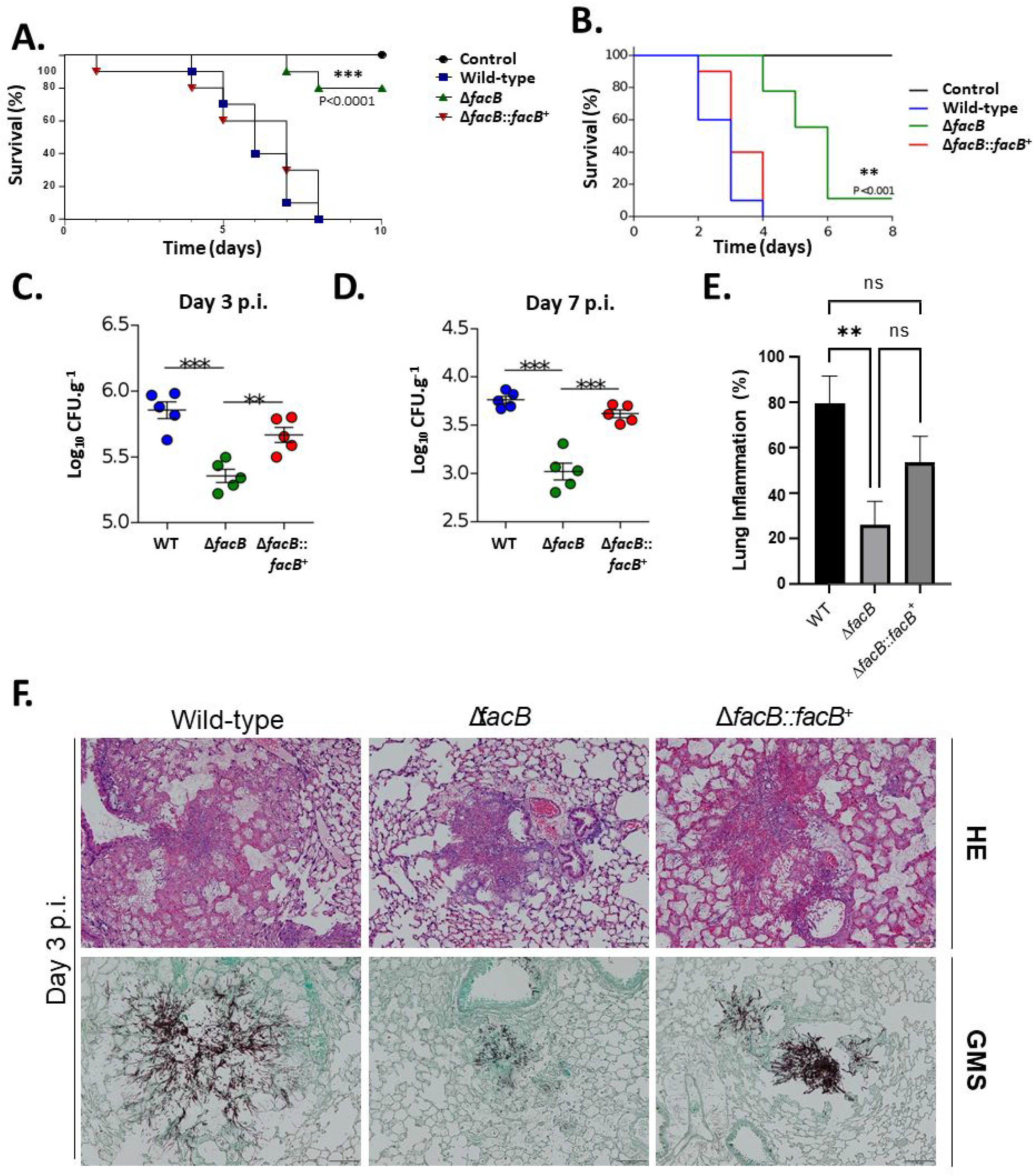
FacB is crucial for virulence in both insect and murine models of invasive aspergillosis. Survival curves (n=10/strain and n=5 for control) of *Galleria melonella* (**A**.) and mice (**B**.) infected with the respective *A. fumigatus* strains. Phosphate buffered saline (PBS) without conidia was given as a negative control. Indicated P-values are based on the Log-rank, Mantel-Cox and Gehan-Breslow-Wilcoxon tests comparing the *facB* deletion strain to the WT and *facB* complemented strains. Fungal burden in murine lungs after 3 (**C**.) and 7 (**D**.) days post-infection (p.i.) with the different *A. fumigatus* strains. Murine lungs were excised, ruptured and re-suspended, before dilutions were prepared that were incubated on plates containing complete medium. Fungal growth was assessed by counting the colony forming units (CFU) on the plates for each dilution. Inflammation in murine lungs after 3 (**E**.) days post-infection (p.i.) with the different *A. fumigatus* strains. Murine lungs were excised and slides of lung sections were prepared. To quantify lung inflammation of infected animals, inflamed areas on slide images were analysed using the thresholding tool in ImageJ software. Standard deviations represent the average of three biological replicates (lungs from different mice) with \**p*-value < 0.01, \*\**p*-value < 0.001, \*\*\**p*-value < 0.0001 in a 2-way multiple comparisons ANOVA test. **G**. Histopathology of mice infected with the different *A. fumigatus* strains. Lungs were excised at 3 days post-infection (p.i.) before lung sections were prepared and stained with HE (Hematoxylin and Eosin) or with Grocott’s methenamine silver (GMS).

To determine whether the observed reduction in virulence of the *facB* deletion strain may be due to growth defects, the WT, Δ*facB* and Δ*facB::facB*^+^ strains were grown for 72 h in the presence of different media that are similar to the mammalian host environment before fungal biomass was quantified. The Δ*facB* strain had significantly reduced growth in the presence of low and high-glucose containing DMEM (Dulbecco’s Modified Eagle’s Medium), FBS (fetal bovine serum) and beef extract when compared to the WT and Δ*facB::facB*^+^ strains but not in the presence of RPMI 1640 medium and minimal medium supplemented with glucose (control) (Supplementary Figure 3E at 10.6084/m9.figshare.14740482). These results suggest that the observed reduction in virulence of the Δ*facB* strain is at least partially due to strain-specific growth defects in a mammalian host environment.

## Discussion

This work aimed at deciphering the regulation of the physiologically relevant carbon source acetate in *A. fumigatus*, and at determining its relevance for fungal virulence. As a first step, we show that *A. fumigatus* can take up and metabolise acetate via the TCA and glyoxylate cycles which is in agreement with studies in *S. cerevisiae* and *A. nidulans* (24, 27). Furthermore, *A. fumigatus* acetate metabolism was shown to be under the regulatory control of the transcription factor FacB, which controls the expression of genes encoding enzymes that are required for the conversion of acetate to acetyl-CoA, for mitochondrial import of acetyl-CoA and enzymes of the glyoxylate cycle. In agreement with the transcriptional data, deletion of *facB* also affected ACS (conversion of acetate to acetyl-CoA) and ICL (glyoxylate cycle) enzyme activities. These results are in agreement with studies in *A. nidulans*, where acetate utilisation as sole carbon source is also dependent on FacB, with this TF regulating the expression of the ACS-encoding gene *facA* and the glyoxylate cycle enzyme-encoding genes *acuD* (ICL) and *acuF* (malate synthase)(25).

In addition, we show that *A. fumigatus* acetate metabolism-related genes as well as ACS and ICL activities are subject to CCR. This is in contrast to findings in *C. albicans* where the addition of glucose to lactate-grown cells did not result in CCR (38). It is important to note here that the experimental conditions differed between our study and (38) (e.g. *A. fumigatus* growth in acetate and glucose versus *C. albicans* growth in lactate then addition of glucose). It is well known that the addition of glucose to cultures causes CCR in *Aspergillus spp* (36), and the rationale here was therefore to present the fungus with equimolar concentrations of both carbon sources during all growth stages. Acetate metabolism was also reported to be subject to CCR in *A. nidulans*, with *facB* and the carnitine acetyltransferase-encoding gene *facC* being under the regulatory control of the CC-repressor CreA (50, 51).

In *A. fumigatus*, CreA-dependent repression of genes encoding enzymes required for acetate metabolism was observed only for the ICL-encoding gene *acuD* whereas the other genes were only partially dependent on CreA-mediated repression. A discrepancy between *acuD* transcript levels and protein activity was observed. It is possible that basal transcript levels are present in all conditions to allow to quickly respond to changes in extracellular available nutrient sources, but that post-transcriptional processing is not taking place. Indeed, gene transcript levels cannot predict protein levels and activity due to mRNA spatiotemporal fluctuations and availability of protein synthesis components (52). Our data suggests that additional repressor proteins and/or mechanisms exist. In agreement, in *A. nidulans*, CreA has been shown to be part of a protein complex that mediates target gene repression, and that co-repressor proteins are crucial for CreA function(53). Furthermore, deletion of *A. fumigatus creA* resulted in significantly decreased expression of acetate utilisation genes in the presence of acetate, suggesting that CreA may be involved in the regulation of these genes in the absence of glucose. In *A. nidulans*, CreA was shown to be important for growth in different carbon, nitrogen and lipid sources and for amino acid metabolism(40); whereas in the filamentous fungus *Trichoderma reesei*, CRE1 was proposed to have roles in chromatin remodelling and developmental processes and was shown to also act as a transcriptional activator (54). These studies suggest that CreA and its homologues have additional roles, other than mitigating CCR, in filamentous fungi.

The deletion of *facB* caused a significantly differential expression of genes that are part of SM biosynthetic gene clusters (BGC) as well as in secreted SMs. A direct correlation between transcript and secreted protein levels is not possible as gene transcript levels cannot predict concentrations of biosynthesised proteins, due to intrinsic fluctuations in mRNA and availability of protein synthesis components (52). Furthermore, SM BGC regulation is extremely complex and governed by many TFs and epigenetic modifications, which result in the expression of different SM BGC in any given condition (55). Subsequently, not all of these SMs are secreted and transcriptional expression of gliotoxin and pyomelanin BGC, as observed here, may be a consequence of the biosynthesis and secretion of other SMs. The role of FacB in the regulation of SM biosynthesis is perhaps not surprising as this TF regulates genes encoding enzymes of central carbon metabolic pathways (e.g. glyoxylate cycle, shown in this work) during growth on alternative carbon sources. SMs are known to be derived from these central metabolic pathways(55) and this study further emphasises that the amount of produced SMs occurs in a carbon source-dependent manner. Alternatively, the observed low concentrations of secreted SMs of the Δ*facB* strains may be due to the inability of this strain to grow in the presence of acetate. Notably, SMs measured in the here defined conditions may be secreted in response to carbon starvation, especially in the presence of low (0.1%) glucose and acetate conditions. These results further emphasise the role of FacB in regulating carbon metabolism. In addition, these results suggest that the concentration of secreted SMs *in vivo* is likely to also depend on host carbon sources and on carbon source starvation, which is encountered within different, nutrient-poor host niches.

Utilisation of different carbon sources also affects the composition of the *A. fumigatus* cell wall, a factor crucial for fungal virulence and pathogenicity and survival within the human host(2). This study shows that growth on acetate results in increased concentrations of the structural polysaccharides β-1,3-glucan and chitin and reduced levels of the cementing, “glue-like” α-1,3-glucan when compared to the *A. fumigatus* cell wall after growth in glucose. Changes in cell wall composition are likely due to differences in primary carbon metabolism that govern the utilisation of these carbon sources and that generate the cell wall polysaccharide precursors(45). Indeed, impairments in *A. fumigatus* glucose utilisation metabolic pathways resulted in an altered cell wall(45). In agreement with other studies, our work suggests that these changes in cell wall composition influence *A. fumigatus* susceptibility to physiological-relevant stresses and antifungal drugs (44, 56). Likely, the significant reduction in the cementing α-1,3-glucan disturbs the organisation of the other cell wall polysaccharides, increasing fungal cell wall permeability and susceptibility to extracellular stresses. This is true for the β-1,3-glucan synthase inhibitor caspofungin and for amphotericin B, which physiochemically interacts with membrane sterols (57). In contrast, increased susceptibility of acetate-grown hyphae to azoles, a class of antifungal drugs that impair ergosterol biosynthesis through targeting lanosterol demethylase of the ergosterol biosynthetic pathway, was not observed. A possible explanation for this may be that caspofungin and amphotericin B both target cell wall and cell membrane components, whereas azoles target intracellular enzymes and that despite the differences in cell wall organisation and polysaccharide content, azole uptake is not affected. In agreement with our data where growth on acetate increases sensitivity to oxidative stress-inducing compounds, *A. fumigatus* acetate-germinated hyphae were more susceptible to human neutrophil-mediated killing *in vitro* when compared to hyphae grown in the presence of glucose. The observed increase in susceptibility to different immune cells is probably due to the differences in cell wall composition resulting from growth in both carbon sources. Our observations are in agreement with a previous study, which showed that a hypoxic environment influenced cell wall thickness, composition and surface exposed polysaccharides, subsequently increasing neutrophil and macrophage reactiveness and activity against *A. fumigatus*(58). The physiological significance of the aforementioned acetate-related differences in cell wall composition, antifungal drug and oxidative stress resistance and interaction with neutrophils remains to be determined. Although acetate was shown to present in the BAL of mice (8), we currently do not know how acetate concentrations fluctuate within certain parts of the lung and the surrounding tissues, as has previously been shown for lung hypoxic microenvironments. It will be interesting to study the distribution of carbon sources in different host niches in future studies.

The Δ*facB* strain exhibited increased susceptibility to macrophage-mediated phagocytosis and killing. Due to the inability of the Δ*facB* strain to grow in the presence of acetate, we were unable to quantify cell wall composition in this strain and therefore determine whether this is a contributing factor when challenged with BMDMs. The inability of the Δ*facB* strain to use acetate may account for the observed increase in phagocytosis and killing of this strain, especially as acetate is available as a carbon source in macrophages(59). In agreement, the expression of genes required for acetate utilisation in the presence of macrophages has been observed for *A. fumigatus* and prokaryotic pathogens(14, 60). It is unlikely that the observed increased phagocytosis and killing of this strain is due to defects in the glyoxylate cycle as previous studies have revealed that glyoxylate cycle enzymes are dispensable for *A. fumigatus* virulence(61),(62). This is further supported by our findings that the utilisation of fatty acids, which results in acetyl-CoA production via β-oxidation(26), is independent of FacB.

Furthermore, the Δ*facB* strain was hypovirulent in both insect and murine models of invasive aspergillosis. Reduced growth of the *facB* deletion strain in media simulating the host environment may account for the reduction in virulence observed for this strain. In addition to acetate metabolism, other FacB-controlled metabolic pathways, which are required for growth in these highly complex nutrient sources may be important for pathogenicity. Investigating the virulence of strains deleted for genes encoding involved in central metabolic pathways such as the phosphoenolpyruvate (PEP) carboxykinase AcuF, the ACS FacA and acetyl-CoA mitochondrial and peroxisome import proteins is subject to future investigations and may further explain the observed reduction in growth. Additional mechanisms may exist, which are important for virulence and are regulated by FacB, especially as the FacB regulon is large. Our RNA-seq data shows that FacB regulates genes encoding proteins important for the production of SMs and oxidoreduction processes, which contribute to virulence. In addition, we cannot rule out that FacB has different targets *in vitro* when compared to *in vivo*, as was previously shown for the *A. fumigatus* AcuM and AcuK transcription factors that are involved in the regulation of gluconeogenesis and iron acquisition(33). The exact mechanism of FacB for *in vivo* virulence thus remains to be determined but is possibly a combination of the aforementioned factors.

In summary, this study describes acetate utilisation in *A. fumigatus* and highlights the importance of carbon source utilisation and metabolic pathways for determining a variety of fungal traits that are crucial for virulence and that potentially shape disease outcome. Future studies should focus on this neglected area of exploring carbon source variety and availability in host primary sites of infection in order to better understand fungal pathogen nutrient requirements and utilisation, which can potentially be targeted for developing anti-fungal strategies.

## Materials and Methods

### Strains and media

All strains used in this study are listed in Supplementary Table 1 at 10.6084/m9.figshare.14740482. Re-introduction of *facB* through homologous combination at the *facB* locus was carried out by co-transformation of the Δ*facB* background strain with the *facB* (amplified by PCR) open reading frame (ORF, no promoter) and the pyrithiamine-containing plasmid pPTR I at a ratio of 2:1. Homologous re-integration of *facB* in the the Δ*facB* locus was confirmed by PCR and by growth assays (Figure 1B). The *facB* deletion mutant was constructed using hygromycin as a selectable marker (32) and is therefore resistant to hygromycin and susceptible to pyrithiamine (Figure 1B). Homologous re-integration of *facB* at the *facB* locus will result in the loss of the hygromycin gene. As pyrithiamine (PT) was used as a marker gene for construction of the re-integration mutant, the resulting strain is resistant to PT (Figure 1B). Growth medium composition was exactly as described previously(40). Radial growth was determined after 5 days whereas dry weight was measured after 3 days of growth. All growth was performed at 37°C and experiments were performed in biological triplicates. Reagents were obtained from Sigma unless otherwise specified.

### Nuclear magnetic resonance (NMR) analysis

Metabolites were extracted from 5 mg freeze-dried fungal mycelia and dried in a speed vacuum as described previously(28). Extracts were reconstituted in 50 μL of deuterated sodium phosphate buffer (100 mM, pH 7.0) containing 0.5 mM TMSP, 3 mM sodium azide and 100% D_2_O. Each sample was sonicated for 10 minutes and vortexed briefly, before a volume of 35 μL was transferred into 1.7 mm NMR tubes.

Spectra were acquired on a Bruker 600 MHz spectrometer equipped with a TCI 1.7mm z-PFG cryogenic probe and a Bruker SampleJet autosampler. One-dimensional (1D) ^1^H NMR spectra and 2D ^1^H-^13^C HSQC spectra were recorded and analysed for each sample as previously described(63).

### RNA extraction and cDNA biosynthesis

RNA was extracted with TriZol (Invitrogen) as described previously(40) and 1 μg of RNA was reverse transcribed to cDNA using the ImPromII^TM^ Reverse Transcriptase kit (Promega), according to manufacturer’s instructions.

### RNA-sequencing

The quality of the RNA was assessed using the Agilent Bioanalyser 2100 (Agilent technologies) with a minimum RNA Integrity Number (RIN) value of 7.0. Illumina sequencing was used for sample RNA-sequencing as described previously(64). Libraries were prepared using the TruSeq®Stranded mRNA LT Set B kit (Illumina) and sequenced (2×100bp) on the LNBR NGS sequencing facility HiSeq 2500 instrument. RNA-sequencing data was processed (quality check, clean-up and removal of rRNA and genome mapping) as described previously(64), with the following modifications. The Bioconductor package tximport (version 1.12.3) was used to import raw read counts into DESeq2 (version 1.24.0), which subsequently quantified differential gene expression. Default Benjamini & Hochberg method was used for multiple hypothesis correction of DESeq2 differentially expressed genes.

### Enzyme activities

Total cellular proteins were extracted as described previously(65) and isocitrate lyase (ICL) activity was measured and calculated as described previously(65). Acetyl-CoA synthetase (ACS) activity was measured and calculated as described previously(66), with the exception that intracellular proteins were extracted as described above and ACS activity was determined in 50 μg total intracellular protein.

### High performance liquid chromatography (HPLC) coupled to tandem mass spectrometry (MS/MS) and data analysis

Fungal biomass was separated from supernatant by miracloth before 20 ml of culture supernatants were freeze-dried. Secondary metabolites (SMs) were extracted from 100 mg freeze-dried sample by re-suspending them in 1 ml HPLC-grade methanol and sonicating them for 1 h in an ultrasonic bath. Samples were filtered and dried under a nitrogen stream before being re-suspended in 1 mL of HPLC-grade methanol. Next, 100 µl of samples were diluted in 900 µl of methanol and passed through 0.22 µm filters into vials.

HPLC MS/MS analysis was performed using a Thermo Scientific QExactive^®^ Hybrid Quadrupole-Orbitrap Mass Spectrometer. Parameters were as follows: positive mode, +3.5 kV capillary voltage; 250 °C capillary temperature; 50 V S-lens and a *m/z* range of 133.40-2000.00. MS/MS was performed using a normalized collision energy (NCE) of 30 eV and 5 precursors per cycle were selected. For the stationary phase the Thermo Scientific Accucore C18 2.6 µm (2.1 mm x 100 mm) column was used. The mobile phase was carried out using 0.1% formic acid (A) and acetonitrile (B) and the following gradient was applied: 0-10 min 5% B up to 98% B; hold for 5 min; 15-16.2 95% B up to 5% B; hold for 8.8 min. The total run time was 25 min and the flow rate was 0.2 mL min^-1^ with 3µL injection volume. Data analysis was conducted using the Xcalibur software, version 3.0.63 (Thermo Fisher Scientific).

Molecular networks were made using the Global Natural Products Social Molecular Networking (GNPS) website (https://ccms-ucsd.github.io/GNPSDocumentation/ from http://gnps.ucsd.edu). First, all MS/MS fragment ions within +/-17 Da of the precursor *m*/*z* were removed and spectra were filtered by choosing only the top 6 fragment ions in the +/-50 Da window for the entire spectrum. The precursor ion mass tolerance and the MS/MS fragment ion tolerance were set to 0.02 Da. Subsequently, networks were created where edges were filtered to have a cosine score higher than 0.6 and more than 5 matched peaks. Edges between two nodes were kept in the network only if each of the nodes appeared in each other’s respective top 10 most similar nodes. Finally, the maximum size of a molecular family was set to 100, and the lowest scoring edges were removed. Network spectra were then searched against the GNPS spectral libraries and library spectra were filtered in the same manner as the input data. Matches between network spectra and library spectra were filtered to have a score higher than 0.6 and at least 5 matching peaks (67). GNPS data used in this work are available at: https://gnps.ucsd.edu/ProteoSAFe/status.jsp?task=f815e5618b05433fb768299 a351fb793 (72 h data).

### Cell wall polysaccharide quantification

Strains were grown for 24 h from 1 x 10^8^ conidia in 50 ml minimal medium (MM) supplemented with 1% (w/v) glucose or sodium acetate. Mycelia were harvested by vacuum filtration, washed, re-suspended in 30 ml of ddH_2_O and disrupted using 5 ml of 0.5 mm glass beads in the FastPrep (MP Biomedicals) homogenizer at 4°C with two cycles of 60 s (6.0 vibration unit) and a 5 min interval between both cycles. Samples were centrifuged at 5000 rpm for 10 min at 4°C, before the cell wall-containing pellets were washed 3 times with ddH_2_O, re-suspended in 15 ml of 50 mM pH 7.5 Tris-HCl, 50 mM EDTA, 2% w/v SDS (2%) and 40 mM β-Mercaptoethanol and boiled twice for 15 min in a water-bath. Samples were centrifuged at 5000 rpm for 10 min and washed 5 times with ddH_2_O. Resultant cell wall fractions were freeze-dried and the dry-weight was measured. Alkali-fractionation of the cell wall was carried out by incubating them twice in 1 M NaOH containing 0.5 M NaBH_4_ at 70°C for 1 h. Samples were centrifuged to separate supernatant [alkali-soluble (AS) fraction] from the pellet [alkali-insoluble (AI) fraction]. The AI fractions was washed six times with ddH_2_O and centrifuged at 5000 rpm for 10 min and freeze-dried. The excess of NaBH_4_ in the alkali-soluble fraction (AS) was neutralized with 2% v/v acetic acid, dialyzed against water until they achieved a neutral pH and freeze-dried. Subsequently, AI and AS fractions were subjected to gas liquid chromatography as previously described(68).

### Minimal inhibitory concentrations (MICs)

MICs of amphotericin B and azoles on the *A. fumigatus* wild-type (WT) strain were carried out as described previously(69) with the exception that the WT strain was also grown in MM supplemented with glucose (GMM) or acetate (AMM).

### Neutrophil-mediated killing of hyphae

Assessing the viability of *A. fumigatus* hyphae in the presence of human neutrophils was carried out as described previously with modifications(70). Briefly, human polymorphonuclear cells (PMNs) were isolated from 8 mL of peripheral blood of adult male healthy volunteers by density centrifugation and re-suspended in Hank’s Balanced Salt Solution (Gibco®). 1 x 10^8^ *A. fumigatus* conidia were incubated for 8 h or 13 h at 37°C in 30 ml GMM or AMM on a rotary shaker before they were centrifuged for 5 min at 4000 rpm, supernatants were discarded and pellets were re-suspended in 1 ml PBS (phosphate buffered saline). To assess the percentage of germinated conidia, samples were viewed under a microscope (Zeiss) at 100x magnification before a total of 100 conidia were counted and the % of germinated conidia was calculated. Pre-grown hyphae were then incubated with neutrophils (0, 1, 2 or 3 × 10^5^ cells / ml) for 1 h at 37°C in RPMI medium before cells were lysed and the MTT [3-(4,5-dimethylthiazol-2-yl)-2,5-diphenyltetrazolium bromide] assay was performed. Hyphal viability was calculated as a percentage of its viability after incubation without neutrophils.

### Bone marrow-derived macrophage (BMDM) phagocytosis and killing assays

BMDM preparation and the ability to kill *A. fumigatus* conidia, as determined by assessing colony forming units (CFU), was carried out exactly as described previously(71). The ability of BMDMs to phagocytise *A. fumigatus* conidia was carried out exactly as described in(72). Fresh *A. fumigatus* conidia were harvested from plates in PBS and filtered through Miracloth (Calbiochem). Conidial suspensions were washed three times with PBS and counted using a hemocytometer. For the killing assay, a dilution of 1×10^5^ conidia in 200 µl RPMI-FCS was prepared. For the phagocytosis assay, 1×10^6^ conidia were re-suspended in 1 ml PBS and inactivated under UV light for 2 h. The percentage of phagocytised conidia was calculated based on conidia cell wall staining with calcofluor white (CFW) (phagocytised conidia are not stained).

### Infection of Galleria mellonella

Breeding and selection of wax moth larvae, preparation of *A. fumigatus* conidia and infection of the last left proleg of larvae with *A. fumigatus* was carried out exactly as described previously(73).

### Ethics statement

Eight-week-old gender-and age-matched C57BL/6 mice were bred under the specific-pathogen-free condition and kept at the Life and Health Sciences Research Institute (ICVS) Animal Facility. Animal experimentation was performed following biosafety level 2 (BSL-2) protocols approved by the Institutional Animal Care and Use Committee (IACUC) of University of Minho, and the ethical and regulatory approvals were consented by the Ethics Subcommittee for Life and Health Sciences (no. 074/016). All procedures followed the EU-adopted regulations (Directive 2010/63/EU) and were conducted according to the guidelines sanctioned by the Portuguese ethics committee for animal experimentation, Direção-Geral de Alimentação e Veterinária (DGAV).

### Infection of chemotherapeutic mice, fungal burden and histopathology

Mice were immunosuppressed intraperitoneally (i.p.) with 200 mg/Kg of cyclophosphamide (Sigma) on days -4, -1, and +2 prior to and post infection, and subcutaneously with 150 mg/Kg hydrocortisone acetate (Acros Organics) on day -1 prior to infection. *A. fumigatus* conidia suspensions were prepared freshly a day prior to infection and washed three times with PBS. The viability of the administered conidia was determined by growing them in serial dilutions on complete (YAG) medium at 37°C. Mice (n = 10/strain) were infected by intranasal instillation of 1×10^6^ conidia in 20 μl of PBS. Mice (n = 5) which received 20 μl of PBS were used as negative control. To avoid bacterial infections, the animals were treated with 50 µg/mL of chloramphenicol in drinking water ad libitum. Animals were weighed daily and sacrificed in case of 20% loss weight, severe ataxia or hypothermia, and other severe complications.

For histological analysis, the lungs were perfused with PBS, excised, and fixed with 10% buffered formalin solution for at least 48 hours, and paraffin embedded. Lung sections were stained with hematoxylin and eosin (H&E) for pathological examination.Paraffin-embedded lung tissue sections were also stained for the presence of fungal structures using the Silver Stain Kit (Sigma-Aldrich), according to the manufacturer’s instructions. Images were acquired using a BX61 microscope (Olympus) and a DP70 high-resolution camera (Olympus). To quantify lung inflammation of infected animals, inflamed areas on slide images were analysed using the thresholding tool in ImageJ software (v1.50i, NIH, USA) according to the manufacturer’s instructions.

### Data Availability

The RNAseq dataset can be accessed at NCBI’s Short Read Archive under the Bioproject ID: PRJNA668271.

## Acknowledgements

We are grateful to the Henry Welcome Building for Biomolecular NMR staff at the University of Birmingham for supporting access to NMR instruments. We would also like to thank the Brazilian Biorenewables National Laboratory (LNBR) for using the NGS sequencing facility to generate the RNA-seq data.

We would like to thank the São Paulo Research Foundation (FAPESP-Fundacao de Amparo a Pesquisa do Estado de Sao Paulo, Brazil) grant numbers 2017/14159-2 (LNAR), 2016/12948-7 (PAC), 2017/08750-0 (TFR), 2016/07870-9 (GHG), 2018/00715-3 (CV) and the Conselho Nacional de Desenvolvimento Científico e Tecnológico, Brazil (CNPq) grant numbers 301058/2019-9 and 404735/2018-5 (GHG) for financial support. IFD acknowledges CICECO-Aveiro Institute of Materials (UIDB/50011/2020 & UIDP/50011/2020) financed by national funds through the Foundation for Science and Technology/MCTES, and the National NMR Network (PTNMR) partially supported by Infrastructure Project N° 022161 (co-financed by FEDER through COMPETE 2020, POCI and PORL and FCT through PIDDAC). AC, RAG, CDO were supported by the Fundação para a Ciência e a Tecnologia (FCT) (PTDC/MED-GEN/28778/2017, UIDB/50026/2020 and UIDP/50026/2020). Additional support was provided by the Northern Portugal Regional Operational Programme (NORTE 2020), under the Portugal 2020 Partnership Agreement, through the European Regional Development Fund (ERDF) (NORTE-01-0145-FEDER-000013 and NORTE-01-0145-FEDER-000023), the European Union’s Horizon 2020 research and innovation programme under grant agreement no. 847507, and the “la Caixa” Foundation (ID 100010434) and FCT under the agreement LCF/PR/HP17/52190003.

Individual support was provided by FCT (SFRH/BD/141127/2018 to CDO, and CEECIND/03628/2017 to AC). This study was financed in part by the Coordenação de Aperfeiçoamento de Pessoal de Nível Superior - Brasil (CAPES) - Finance Code 001 (JHC scholarship). SSWW was supported by Pasteur-Roux-Cantarini fellowship. JLS and AR are supported by the Howard Hughes Medical Institute through the James H. Gilliam Fellowships for Advanced Study program; AR is additionally supported by the National Institutes of Health / National Institute of Allergy and Infectious Diseases (1R56AI146096-01A1).

